# Brain activation for language and its relationship to cognitive and linguistic measures

**DOI:** 10.1101/2024.12.31.630909

**Authors:** Irene Balboni, Alessandra Rampinini, Olga Kepinska, Raphael Berthele, Narly Golestani

**Affiliations:** Department of Psychology, Faculty of Psychology and Education Science, University of Geneva, Geneva, Switzerland; Vienna Cognitive Science Hub, University of Vienna, Vienna, Austria; Department of Behavioural and Cognitive Biology, Faculty of Life Sciences, University of Vienna, Vienna, Austria; Institute of Multilingualism, University of Fribourg, Fribourg, Switzerland

**Keywords:** Language, Cognition, Brain-Behaviour Relationships, Dyslexia, Bilingualism

## Abstract

Language learning and use relies on both domain-specific and domain-general cognitive and sensory-motor functions. Evolutionary and developmental perspectives, as well as the use of language in interaction, suggest an overlap between language and other skills. Building on our previous behavioural findings, which outlined a consistent behavioural association between language and domain-general skills, we investigated brain-behavioural associations using story listening during fMRI and behavioural measures of language, reading, multilingual experience, cognition, musicality, arithmetic, and motor skills. Participants varied in multilingual language experience and reading aptitudes, including both typical (TRs) and dyslexic readers (DRs). Using multivariate Partial Least Squares correlation, we identified a main component linking cognitive, linguistic, and phonological measures to brain areas underlying lexico/semantics, combinatorial processing, and amodal semantic processing. A second analysis excluding DRs showed closer associations between cognitive/linguistic, literacy, phonological and memory processes within the same brain network as in the full sample. Here, we also isolated additional, complementary components, including one involving speed, automatization and lexical access, linked to auditory and motor brain areas. This suggests greater coherence and more integrated, ‘expert’ processing in TRs. This work is a first step in exploring complex relationships between language and non-linguistic functions that are important to it.

## 1. INTRODUCTION

Language is a key human ability and is represented by a complex system of spoken, signed or written symbols allowing us to communicate with others (Crystal and Robins 2024 Jul 25). It enables abstraction and is believed to support complex thinking and cognition (Carruthers 2002; Majid et al. 2004). Language is mostly learned and used during face-to-face interactions, and with the support of paralinguistic and non-verbal cues (Levinson and Holler 2014). Being a successful language user entails a remarkable number of processes and subroutines, and yet neurotypical individuals successfully learn and use a first (L1) and often a second (L2) language. The most established models of the neurobiology of language perception and production highlight the hierarchical nature of language, involving phonological, lexico-semantic and morphosyntactic information, subserved neurally by a left-lateralised perisylvian network including primarily dorsal motor/premotor areas and inferior frontal areas, and superior, middle and inferior temporal cortices (Hickok and Poeppel 2004; Hickok and Poeppel 2007; Rauschecker and Scott 2009; Rauschecker 2012; Rauschecker and Scott 2016). Some authors argue for a broader inclusion of areas in the so-called ‘language network’, like the inferior parietal cortex, underlying knowledge representation, as well as the anterior cingulate and dorsolateral prefrontal cortices for control functions (Price 2012; Hagoort 2014; Hagoort 2017; Hagoort and Özyürek 2024), while others take a conservative approach and consider only areas strictly involved in the access to meaning and sentence construction, including only left-lateralized lateral frontal (IFG, middle frontal) and superior temporal areas (Fedorenko et al. 2024).

Language is a uniquely human and relatively recent evolutionary skill, especially in the case of literacy. To understand how our brains evolved to accommodate such a new and complex skill, different theories and findings suggest that specific aspects of language had to rely on pre-existing mechanisms that predated the emergence of language and that could be used and specialised for language because of their computational properties (Asano and Boeckx 2015; Asano et al. 2022). This has been described as neural reuse or ‘exaptation’ (Fitch 2011; Fitch 2012; Janacsek et al. 2022), and may explain why language has often been associated with other domain-general and domain-specific abilities and why the question of language specificity is difficult to resolve.

Different implementations could underlie this overlap between language-specific and other domain-general or domain-specific functions. One possibility is that different abilities might leverage the computational mechanisms of a specific brain area. The computational mechanisms themselves may be shared across domains, while domain-specificity may be shaped by the networks in which it is embedded (Asano and Boeckx 2015; Asano et al. 2022). This has been proposed to be the case for the IFG, specifically BA44/45, which is known to be involved in syntax processing in language, but is thought to have evolved for this due to its role in hierarchical motor planning (Asano and Boeckx 2015; Asano et al. 2022). As such, this region is thought to commonly underlie hierarchical processing for music, action, and language (Fitch 2011; Fitch and Martins 2014; Asano et al. 2021; Asano et al. 2022). A second possible mechanism that has been proposed to explain the neural underpinnings of language in brain regions originally involved in a different function is that of a complete replacement of old functions with new ones. A widely accepted example of this ‘neuronal recycling’ comes from research on developmental brain adaptations to reading, a late-emerging skill both in evolutionary and developmental terms. Initially involved in face and object perception in preliterate children, this area is repurposed for reading after acquisition. As a result, in adults, this area responds reliably to words as an integral part of the language system, while its sensitivity to faces and objects decreases (Cohen and Dehaene 2004; Dehaene 2005; Dehaene et al. 2015; Dehaene-Lambertz et al. 2018). These different explanations for shared mechanisms between language and other domain-general (e.g., cognition, perception, motor skills) and domain-specific skills (e.g., musicality) processing can explain observed neural overlap across language and other domains.

Beyond brain regions that may have originally served different purposes in ontogeny or phylogeny, naturalistic use of language – especially under more demanding conditions – involves interaction with other cognitive functions (executive control, memory) and modulation from bottom-up sensory areas. This has been suggested by ‘declarative-procedural’ memory model of language, one of the most prominent language models (Ullman 2004; Ullman 2016). The model posits that the native language acquisition of lexicon and grammar depends on general-cognitive mechanisms of declarative and procedural memory, respectively, with their associated neural networks playing a crucial role in both language functions and general memory abilities. Moreover, given that meaning interpretation is highly context-dependent (Hagoort and Van Berkum 2007) and the processing of the speech input is influenced by top-down information (McClelland et al. 2006; Davis and Johnsrude 2007; Rutten et al. 2019), it is more plausible that language processing engages broader cognitive mechanisms. It is also important to consider that the use of language in successful communication (i.e., in interaction with others) goes beyond the single word processing paradigm often used in neuroimaging studies, and it is likely to rely on additional cognitive processes, such as attention or theory of mind in order to deal with the demands like turn-taking and tracking the narrative of the conversation (Hagoort 2014). Lastly, given the multimodal nature of language, which requires the integration of oral, written and also paralinguistic information, (Levinson and Holler 2014; Cohn and Schilperoord 2024; Hagoort and Özyürek 2024), effective processing is likely to depend on the coordination of wide multimodal cortical areas and binding of top-down and bottom-up information, likely to be enabled by subcortical structures, such as the basal ganglia, thalamus and cerebellum (Asano et al. 2022; Janacsek et al. 2022; Turker et al. 2023). Taken together, this literature suggests that the neural substrate supporting language may extend beyond traditional models that include only perisylvian cortical regions and has led some researchers to view language as “widely distributed throughout the brain” (Aliko et al. 2023; Drijvers et al. 2025).

At the behavioural level, and in particular in the study of language aptitude (i.e., the potential for language learning), various models and theories already consider skills beyond language-specific abilities necessary to make a good language learner. Language aptitude has been linked to musical abilities (Nardo and Reiterer 2009; Fonseca-Mora et al. 2011; Turker et al. 2017; Turker and Reiterer 2021; Coumel et al. 2023), as well as general cognition (Doughty et al. 2010) with particular attention to the role of memory (Wen 2012; Singleton 2017; Feng et al. 2021). Also, in behavioural studies L1 processing language-specific abilities have been consistently found to interact with general cognition during development (for a review: Kidd & Donnelly, 2020) and to be supported by cognition in healthy older adults to ensure language comprehension despite sensory and cognitive decline (Wingfield and Grossman 2006), particularly during the perception of speech in noise (Wong et al. 2010). Moreover, language is associated with early motor abilities (Alcock and Krawczyk 2010; Libertus and Violi 2016; Gonzalez et al. 2019), as well as music (for an overview: Nayak et al., 2022) and numerical skills (Simmons and Singleton 2008; Purpura and Lonigan 2013). It is worth noting that while these skills are positively linked to language abilities in individuals with typical development, they also show deficits in dyslexia, a neurodevelopmental disorder that leads to poor literacy outcomes (Lyon et al. 2003). Dyslexia has been associated with impairments in motor control (Marchand-Krynski et al. 2017; Decarli et al. 2024) and musicality (Overy et al. 2003; Christiner et al. 2022), with musical training being explored as a potential remediation therapy (Overy 2003; Rolka and Silverman 2015). Additionally, difficulties in numerical skills are common, as dyslexia and dyscalculia frequently co-occur and share genetic influences (van Bergen et al. 2023)

In order to learn about the componential structure of language and cognition and of the relationships therein, individual differences can be leveraged to explore which abilities stably associate and dissociate in light of the variability (Bates et al. 1991; Kidd et al. 2018). Several behavioural studies having assessed cognitive and linguistic performance used this approach to explore the complex interplay between these domains, and showed in adults (Hintz, McQueen, et al. 2024; Hintz, Voeten, et al. 2024) and in children (Berthele and Udry 2021; Udry and Berthele 2021) that language-specific and domain-general cognition cannot be clearly separated. In light of these works and of the above-described debate, in previous work we collected data from a relatively large participant sample (*n* = 152) on behavioural measures of language, cognition, musicality, arithmetic and motor skills, as well as on measures of language and reading experience to explore patterns of dissociation and association between these domains (Rampinini et al. 2024). In order to maximise inter-individual variability, we recruited people with a wide range of multilingual language experience, and we also included a subgroup of adults who had previously been diagnosed with dyslexia. Using exploratory graph analysis, we analysed the network of relationships between individual variables while identifying underlying clusters. We confirmed that language and cognition are stably associated, and that none of the identified clusters were exclusively composed of language-specific or of domain-general cognitive skills. On the other hand, musicality, a domain-specific skill often associated with language, was isolated from language and cognition scores. The same pattern of isolation was present for multilingualism measures and literacy, indicating that these measures are possibly more linked to experience and education and that they are dissociated from performance in language/cognition.

Less work has explored the key dimensions supporting language processing beyond language-specific skills and their corresponding neural correlates in a multivariate manner. Although recent work has started to highlight the importance of multimodal and multivariate analyses of language learning profiles, this approach remains relatively underexplored (Feng et al. 2021). In the present work, we build upon our previous results at the behavioural level to extend our study to brain activation for native language speech processing (story listening). L1 speech processing is an informative and stable neural marker (Mahowald and Fedorenko 2016) of how language is represented in the brain and of individual differences in language processing and experience. Neural responses to L1 have been linked to outcomes in L2 learning (Díaz et al. 2022; Fuhrmeister et al. 2023), multilingual experience (Jouravlev et al. 2020), and print processing that underlies successful language reading (i.e. print-speech convergence; Rueckl et al. 2015). Furthermore, the use of data collected during L1 as opposed to L2 speech listening allows individual variation to be sampled while maintaining a degree of homogeneity in terms of age of acquisition and baseline level of exposure. We included whole-brain activation, including subcortical areas, to assess brain-behaviour associations in a fully data-driven manner. This allowed us to avoid limiting the analysis to traditional language areas and implicitly favouring one ‘language model’ over another. To uncover this relationship, we used Partial Least Squares Correlation (PLS), a method that identifies components explaining common variance between two types of data.

## 2. METHODS

### 2.1. Stimuli and procedure

#### 2.1.1. Participants

152 participants took part in the study. Participants were healthy, mostly speaking French as a first language and having knowledge of two or more languages. None of the participants had official qualifications for language translation or interpretation, nor did they have professional musical training. Participants who spoke more than 6 languages were included in the study even if French was not their native language, if they reported advanced French proficiency. Figure 1 depicts the multilingual background of the participants in the sample included in the analysis, TRs had on average higher multilingualism scores than DRs (as shown also in the Supplementary results). Details of the participants’ language and demographic backgrounds are available in the supplementary materials (Table 1). The final sample only included participants who did not have missing data on any of the measures analysed. It consisted of 134 participants, aged between 18.1 and 47.2 years (*M* = 24.0, *SD* = 4.9, 92 females). A subset of these participants *(n =* 25) was diagnosed with dyslexia before taking part in the study. A second analysis was run only on 109 typically reading participants (TRs), again selected based on having no missing data. In this subsample, participants were between 18.2 and 47.2 years of age (*M* = 24.1, *SD* = 5.07, 74 females).

**Figure 1.**
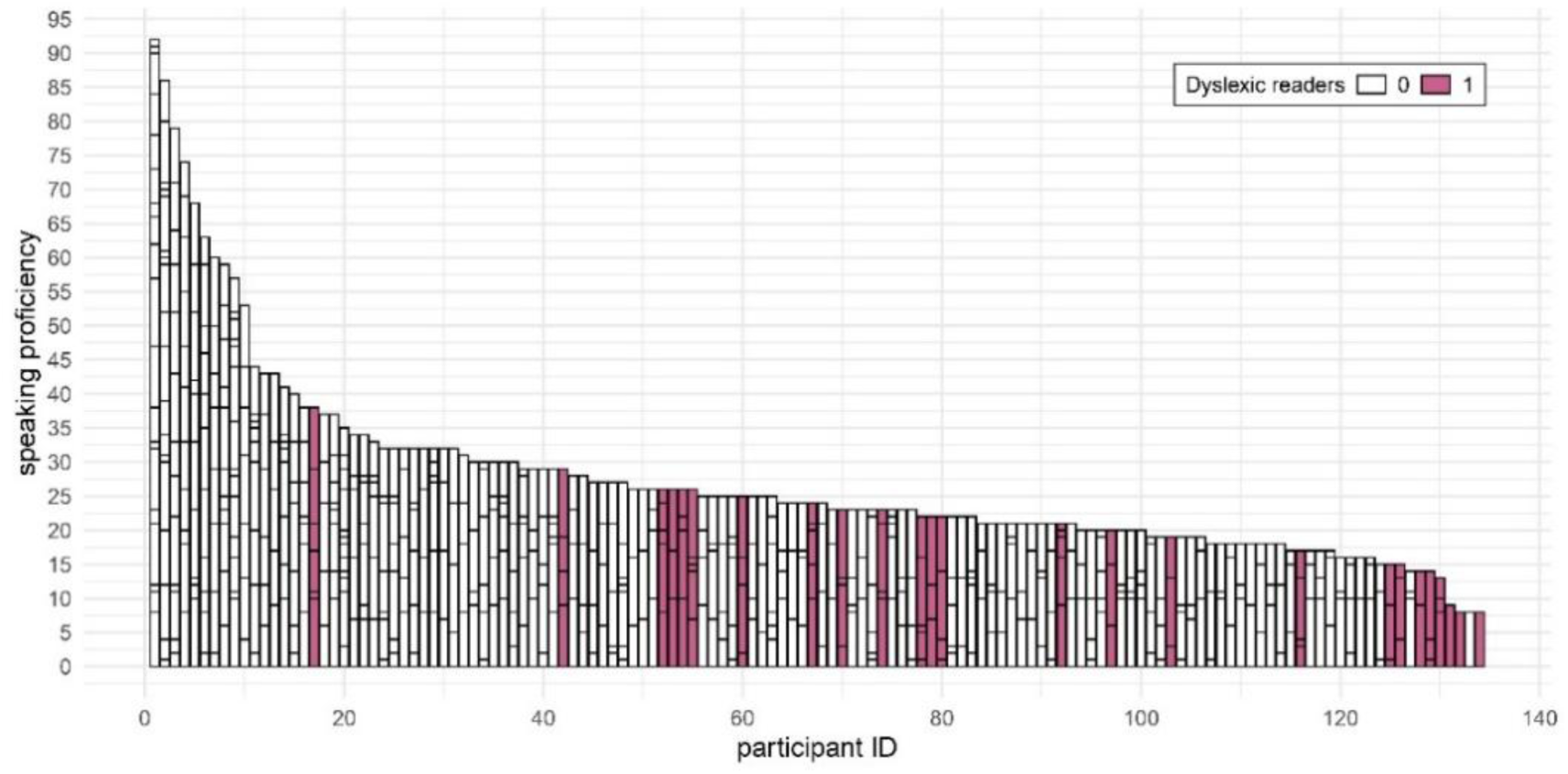
Language background of the participants in the sample. Each bar represents the language experience of one participant; the height of the stacked bars within each bar represents the self-reported speaking proficiency for individual languages (the taller the bar, the higher the speaking proficiency for that language). Before plotting, the data was sorted based on cumulative language experience values; as a result, the left-hand side of the figure displays hyperpolyglots’ data while the right-hand side shows data from monolinguals. Dyslexic readers are marked in dark pink.

**Table 1.**
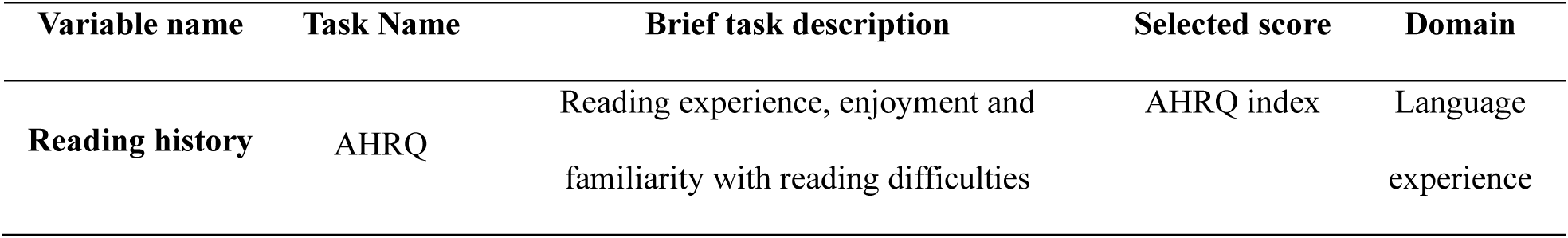
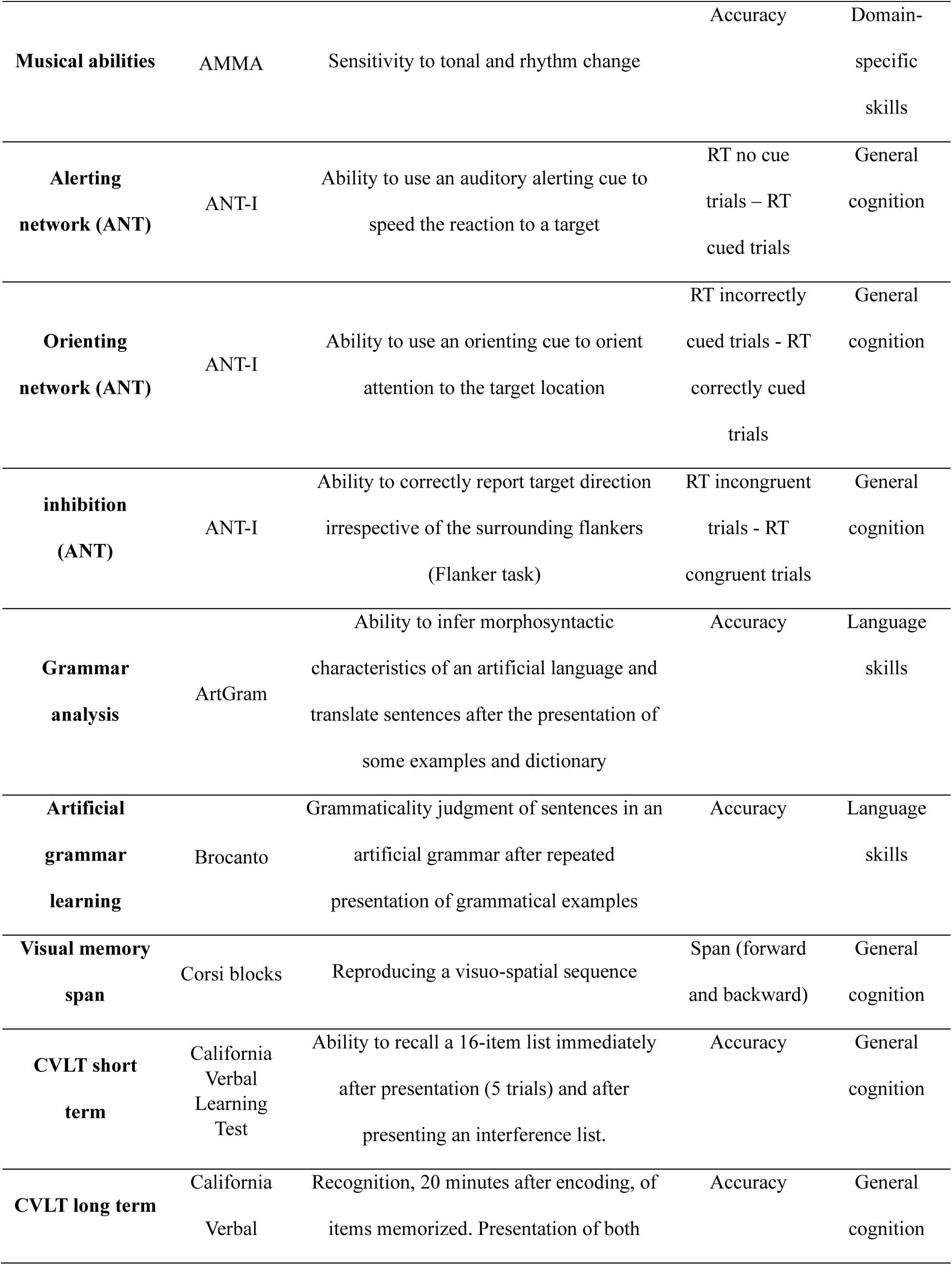
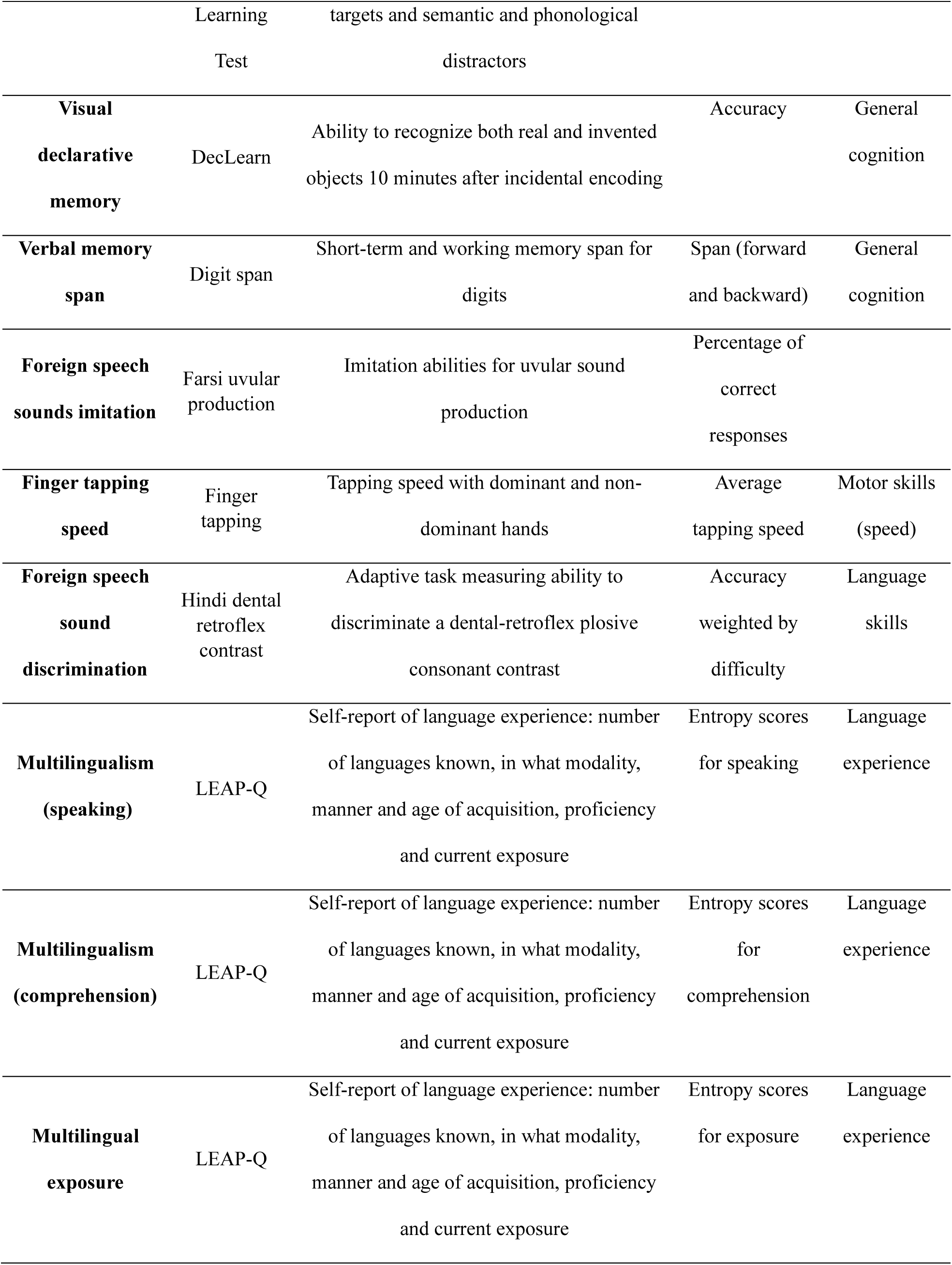
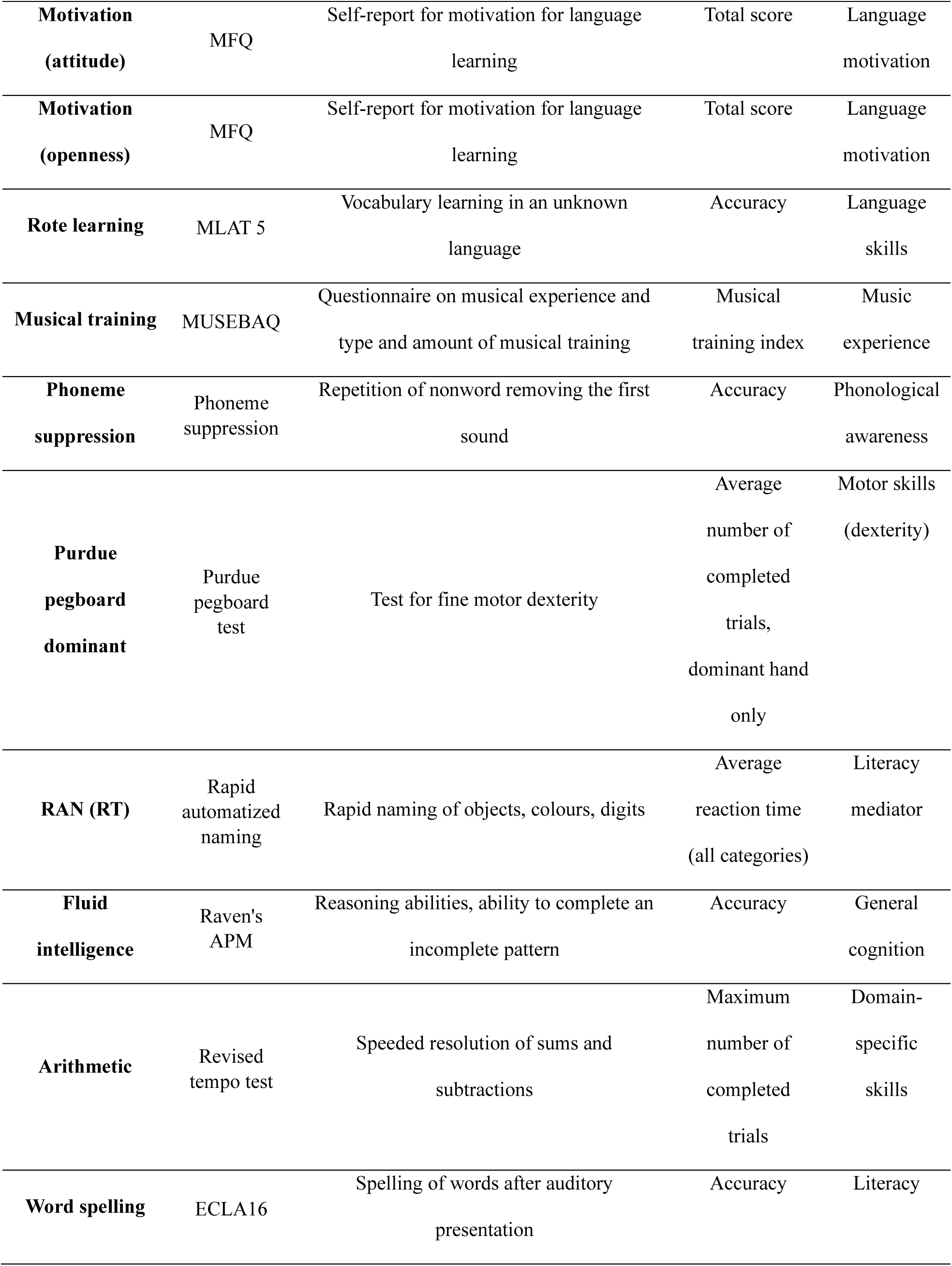
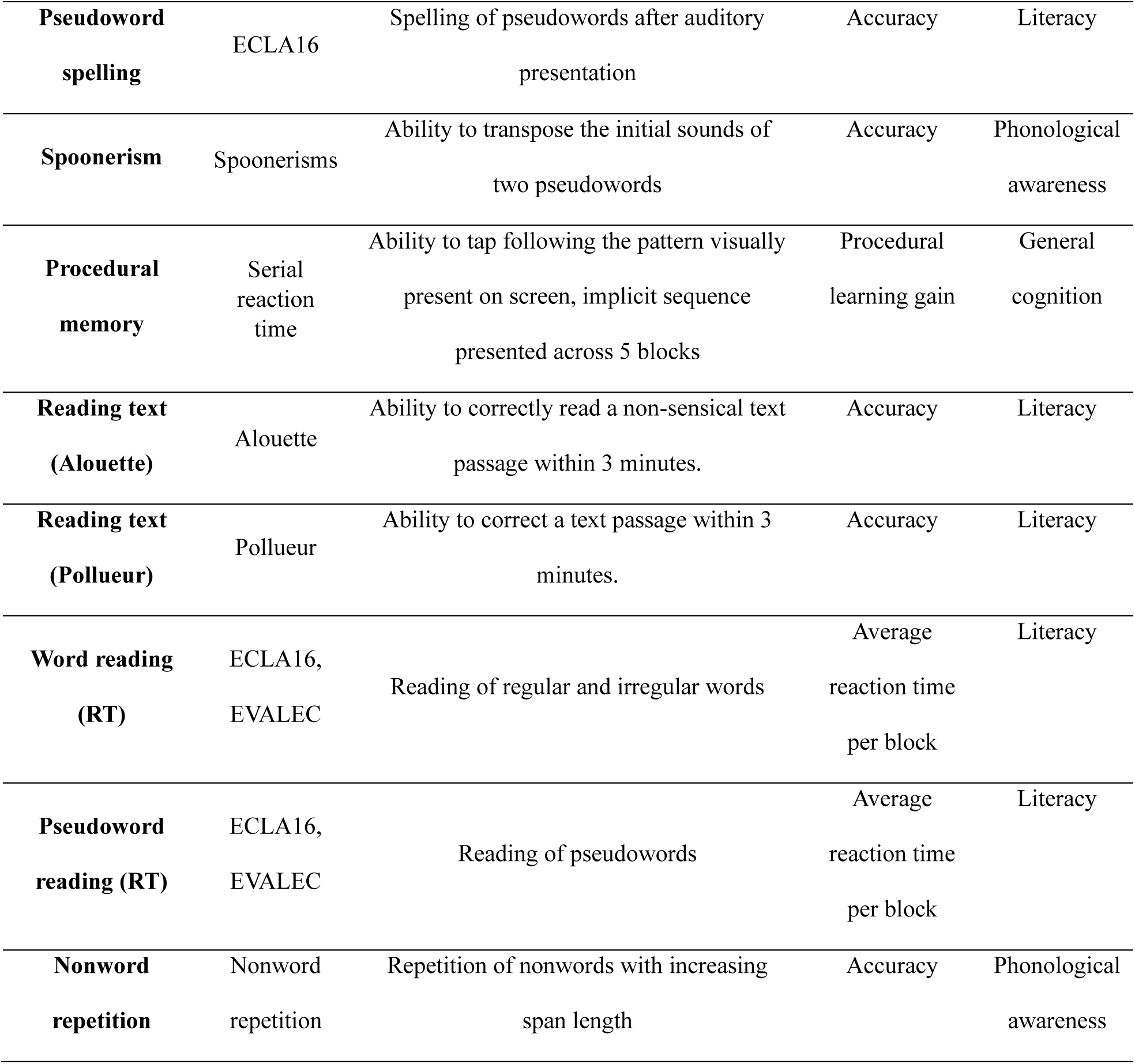
Overview of the tasks, scores and resulting variables obtained from behavioural testing. All the scores, including reaction times and other metrics used in this analysis, were transformed such that higher values consistently indicate better performance.

#### 2.1.2. Procedure

Each participant took part in 4 different sessions (spread over several testing days): session 1 to fill in online questionnaires, session 2 to perform a portion of the tasks online while supervised by one trained member of the team, session 3 to complete in-person tasks that required closer supervision or standardised hardware and session 4 for neuroimaging and genetic data collection (the latter not included in this study). Questionnaires (session 1) were delivered using Qualtrics XM©, online tasks (sessions 2 and 3) were delivered using Gorilla, and the fMRI task (session 4) was presented using Psychtoolbox3 (Matlab r2021b). An overview of all the tasks and behavioural measures is presented in Table 1. Detailed description of the hardware, procedure, and extended descriptions of the behavioural tasks and their references can be found in our previous work (Rampinini et al. 2024). All eligible participants provided signed informed consent for all subsequent experimental procedures and data reuse. The study received ethical approval from the Geneva Cantonal Ethical Commission (Protocol N. 2021-01004).

#### 2.1.3. fMRI data collection

All participants took part in an MRI session where we collected brain structural (MPRAGE), MP2RAGE, DWI) data, and functional imaging data during resting-state, and during music, reading and language localizers, the latter both in the participants’ L1 and L2. In this work, we only analysed fMRI data from the language localiser in the L1, and used the T1 MPRAGE sequences for co-registration. Images were acquired using a Siemens 3T Magnetom-Prisma scanner at Campus Biotech (Geneva), equipped with a 64-channel head coil. T1-weighted MPRAGE structural images were collected using 208 sagittal slices (TRs = 2,300 ms, TE = 3,26 ms, voxel size: 1×1×1 mm;). BOLD activation was measured using a whole-brain EPI sequence (acceleration factor −1, FOV=224 mm, voxel size: 2×2×2 mm, TRs = 2000,0 ms, TE=32,00 ms flip angle 75°, interleaved acquisition, 72 slices).

During the fMRI task, participants passively listened to intact and degraded speech passages of the story ‘Alice in Wonderland’ while fixating a black fixation cross on a white screen. The stimuli and task are openly available (http://web.mit.edu/evlab/aliceloc/index.html), and the procedure for the degradation of low-level auditory features is described in Malik-Moraleda and colleagues (2022). We used the localiser to present intact and degraded passages for both the L1 and the L2 to each participant. In total, 3 runs were acquired, each including 5 fixations blocks of 12 seconds and 16 blocks of 18 seconds each, 4 per each of the intact and degraded conditions. No response was required, and participants took short, self-timed breaks between blocks. Of the 134 participants included in the final analysis, a total of 127 listened to passages in French for the L1, while the remaining 7 chose another language (this always either being their first language or a language that felt native to them): 3 participants chose English, 1 chose Spanish, 1 chose German and 1 chose Portuguese. More information on the individual profiles of each participant can be found in the supplementary material (Table 1).

### 2.2. Analysis

#### 2.2.1. Behavioural data

Behavioural data for the questionnaires and the tasks completed on the Gorilla online platform were analysed and scored using in-house Python scripts. Tasks requiring manual scoring, such as spelling or foreign speech sound imitation, were scored manually by native speakers. All measures, with their corresponding tasks and scores, are outlined in Table 1. A more detailed description of the behavioural tasks, together with descriptive results and reliability indices for each task, can be found in Rampinini and colleagues (2024). The supplementary materials (Figure 1) include plots with scores for all participants in all of the included behavioural measures, divided by reading group.

#### 2.2.2. Preprocessing fMRI data

Functional images were preprocessed using FSL 6.0. Preprocessing was carried out using FEAT (FMRI Expert Analysis Tool, version 6.0) with a standard preprocessing pipeline that included the following steps: removal of non-brain tissue using optiBET (Lutkenhoff et al. 2014), B0 unwarping using fieldmap images and nonlinear registration to structural T1 images (BBR, FNIRT), and spatial smoothing using a Gaussian kernel of 5 mm FWHM. Data were motion-corrected using ICA-AROMA (Pruim et al. 2015) and no participant had an absolute displacement higher than 5 mm (which would have resulted in their exclusion from the sample). We then removed nuisance regressors (white matter, WM, and cerebral-spinal fluid, CSF), performed high-pass filtering at 90 s and registered the preprocessed data to the MNI152 standard space (Grabner et al. 2006). The BOLD response was modelled using a double-gamma haemodynamic response function (HRF). Activation for intact and degraded L1 speech listening was modelled, and the intact *vs.* degraded speech contrast was generated.

#### 2.2.3. Extraction of individual level activity with parcellation

Once the intact *vs.* degraded brain activation contrast was computed for each participant, we defined different brain parcels in order to maintain a whole-brain approach while reducing the numbers of brain variables for the PLS analysis (i.e. as compared to a voxel-based approach). We included all cortical and subcortical regions. For the cortical regions, a total of 397 parcels, or regions of interest (ROIs) were identified. The cortex was segmented using the Glasser multimodal parcellation (Glasser et al. 2016). The parcellation was adapted to volume space, registering it to the same MNI152 template as the functional data (Horn 2016 Jun). The volume was then split by hemisphere to obtain 180 parcels per hemisphere. In the results section, cortical ROI labels correspond to each area’s name in the original parcellation publication (Glasser et al. 2016). ROIs for the hippocampus were excluded from the cortical parcellation because hippocampal areas were defined using a subcortical atlas, as follows: subcortical regions comprising bilateral hippocampus, thalamus, amygdala, pallidum, putamen, and caudate nucleus were identified using the Harvard-Oxford subcortical atlas (Makris et al. 2006). Cerebellar regions were identified using a probabilistic cerebellar atlas (Diedrichsen et al., 2009; full list of cerebellar ROIs in the supplementary material). Both subcortical and cerebellar areas were extracted using a threshold of 50%. The full list of ROIs is presented in the supplementary materials. For each of the ROIs obtained using the above parcellations, average contrast values for the difference in brain activation for intact *vs.* degraded speech were extracted. The resulting brain dataset extracted for each participant contained the average contrast values for each of the 397 ROIs.

#### 2.2.4. PLS

We used PLS to evaluate the multivariate relationships between two data modalities, brain and behavioural, aiming to isolate components that maximise shared variance between the two sets of data. PLS is conceptually similar to principal component analysis, which finds patterns within a single dataset, with the different that PLS extracts components that best explain the correlation between two sets of variables. PLS works by computing linear combinations of variables from both datasets to uncover hidden relationships. This makes it a powerful tool for studying complex interactions between different types of data.

We implemented it by first z-scoring both datasets and correcting them for age, sex and handedness through residual calculation (McIntosh and Lobaugh 2004; Krishnan et al. 2010; Krishnan et al. 2011). After data preprocessing and preparation, the analysis was performed using the myPLS toolbox (https://github.com/danizoeller/myPLS). A correlation matrix between behavioural and brain data was generated, and then the main components which explained the relationship between the two modalities were extracted through singular value decomposition.

For each component, behavioural and brain weights were calculated: these explain how strongly each single variable contributed to the multivariate correlation between the data modalities in that component. Behavioural weights, or saliences, represent how strongly each variable contributes to the correlation between brain and behaviour in each component. Brain weights, or saliences, represent how strongly the intact versus degraded speech listening average contrast values within each ROI contributes to the correlation between brain and behaviour in each component.

The significance of each component was assessed using 1000 permutations, and the stability of the weights of each variable per component was evaluated using a bootstrapping procedure (500 samples). The significance of each variable was determined using a *Z* score >=|3| threshold, which corresponds to a two-tailed probability of 0.0013. Previous work suggested that a Z score >=|2| (Krishnan et al. 2011) or >=|2.57| (equivalent to a two-tailed probability of 0.01; McIntosh & Lobaugh, 2004) could be used, but in the present analysis, given the high number of variables used in both datasets, we chose a stricter threshold to consider the results robust/significant. The results report weight means and standard deviation after bootstrapping for each variable/ROI. More details on permutation, bootstrapping and thresholding procedures are available in the Supplementary Materials (Supplementary methods section).

As mentioned in the Participant section, we initially ran the analysis on the full sample including TRs and DRs (*n* =134), and then on a subsample including TR participants only (*n*=109). We could not perform a separate analysis on the DRs alone due to their smaller sample size (*n* =25), which, given the number of variables included in the analysis, would likely have resulted in overfitting and poor generalization (Mihalik et al. 2022). To ensure that any potential differences between the two analyses were due to sample characteristics and not to differences in sample size, we ran an additional PLS analysis on a third subsample, hereafter referred to as the verification subsample.

This had the same sample size as the subsample of TRs only (n=109), but included the same proportion of TRs and DRs as the full sample.

## 3. RESULTS

### 3.1. Partial least square results

Overall, the analysis of the whole sample resulted in one significant component whereas the analysis including only TR participants resulted in 3 significant components. The analysis from this verification subsample only involving TR participants showed a substantial overlap with the results in the whole sample, confirming that differences between the latter and the subsample were likely due to differences driven by sample characteristics and not sample size reduction. All the results for the verification subsample are available in the supplementary material.

#### 3.1.1. Brain and behavioural relationships in the whole sample

We applied a PLS analysis to the 36 behavioural measures and to the average activation (contrast values for intact *vs.* degraded L1 speech) from the 397 cortical and subcortical areas during L1 story listening. One component was significant after permutation testing (*p* = .006, *r* = .34). Figure 2 shows the contribution of each variable from both modalities to the brain-behavioural correlation component (more details are available in the Results section of the Supplementary Material). The component explained 36% of the brain-behavioural covariance.

**Figure 2.**
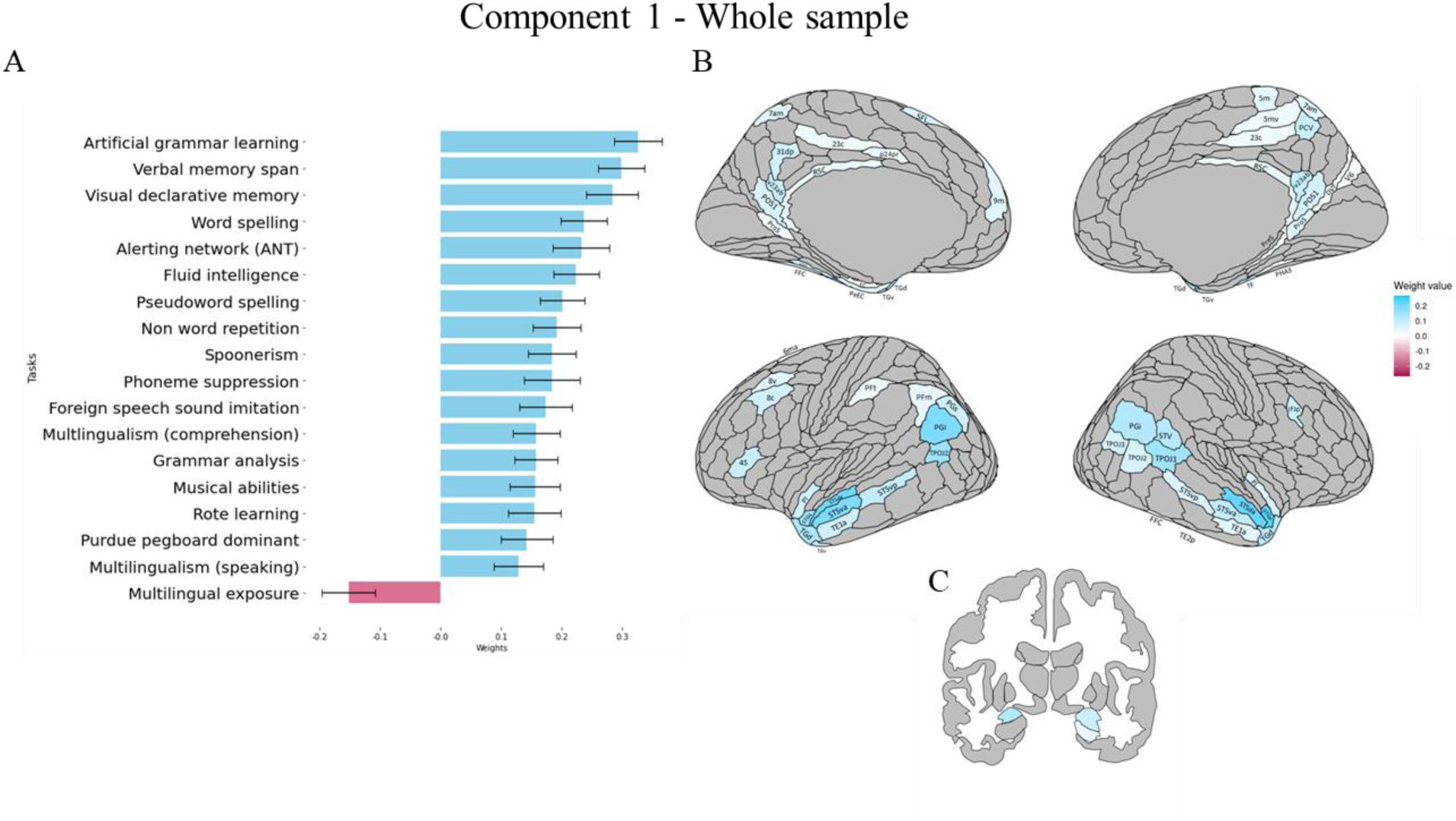
Results of the behavioural, cortical and subcortical saliences/weights for the significant correlational PLS component in the whole sample. Blue represents significant positive weights, red represents significant negative weights, and grey represent non-significant ones. (A) Contribution of behavioural variables to the component (for all scores, higher values represent better performance). (B) Contribution of cortical ROIs to the component. (C) Contribution of subcortical areas to the component. For each component, mean weights, SD and brain uncorrected results can be found in the supplementary material, in the Supplementary results section.

At the behavioural level, the component was mostly positively associated with half of the behavioural tasks (Figure 2A). This component identified an association between measures of general cognition (memory in particular), language skills (with a prominent role of grammar) and to a lesser extent of phonological skills. More specifically, a positive association was found between the brain measures and measures of language and cognition that rely heavily on memory processes (such as nonword repetition, declarative and verbal memory, and rote learning), pattern recognition and learning (morphosyntactic patterns for a grammar analysis task, and visual patterns for fluid intelligence scores), or a combination of both (as seen in artificial grammar learning). Artificial grammar learning skills were, both in the full sample and in the typical readers only (see next analysis) the strongest contributors to the component. Another set of measures pertain to the auditory domain and to auditory working memory: musicality and phonological awareness (including spoonerisms, nonword repetition, and foreign speech sound imitation), and to a literacy measure involving sound-symbol correspondences (i.e. word spelling). Multilingualism scores, with a stronger influence of multilingual comprehension than of speaking, also contribute positively to this component. Moreover, there was a positive contribution of alerting skills, i.e. a measure of response readiness and of sustained attention to incoming stimuli (Ishigami et al., 2013), and of motor dexterity (Purdue pegboard test). Last, Multilingual exposure, differently from multilingual comprehension and speaking abilities, contributed negatively to the component, with a relatively low weight compared to positively associated variables.

All cortical and subcortical brain regions showing a significant contribution to the component, and their weights, are shown in Figure 1B (results for cerebellar ROIs and uncorrected results can be found in the Supplementary Materials). Overall, better performance on the above tasks correlated with stronger activation of bilateral anterior temporal cortical areas (e.g. STGa, STSda, STSva, TGd, TGv, TE1), posterior cingulate areas (e.g. RSC, v23abm, POS1), inferior parietal regions (PGi, PGs, PFm), and temporo-parietal-junction (TPJ; TPOJ1, TPOJ2, TPOJ3). The latter regions – inferior parietal cortex and adjacent temporo-parietal junction – showed somewhat left *versus* right hemisphere lateralization, respectively. Other areas that displayed a significant positive activation in relation to behavioural measures comprised predominantly left-lateralised dorsolateral prefrontal (8v, 8c) and inferior frontal (IFJp, 45) regions. Better scores also positively correlated with activity in medial temporal regions such as the entorhinal cortex (EC) and perirhinal cortex (PeEC) in the left hemisphere, presubiculum (PreS) in the right hemisphere and para-hippocampal (PHA3) areas bilaterally. Subcortically, Figure 2c shows a positive correlation with bilateral amygdala and right hippocampus. None of the cerebellar regions were significantly associated with the component.

#### 3.1.2. Brain-Behavioural relationship in typical readers only

We applied a second PLS analysis to the same behavioural and brain measures presented in the above section, but this time, we included only participants without a dyslexia diagnosis *(n* = 109). After permutation testing, we identified 3 significant components.

##### Component 1

The first component accounted for 44% of the variance (*p* = .001, *r* = .45). Behaviourally, this component resembles the one identified in the whole sample, but it is associated with a higher number of memory measures and, differently from the whole-sample analysis, includes a larger number of literacy measures (brain and behavioural saliences are shown in Figure 3).

**Figure 3.**
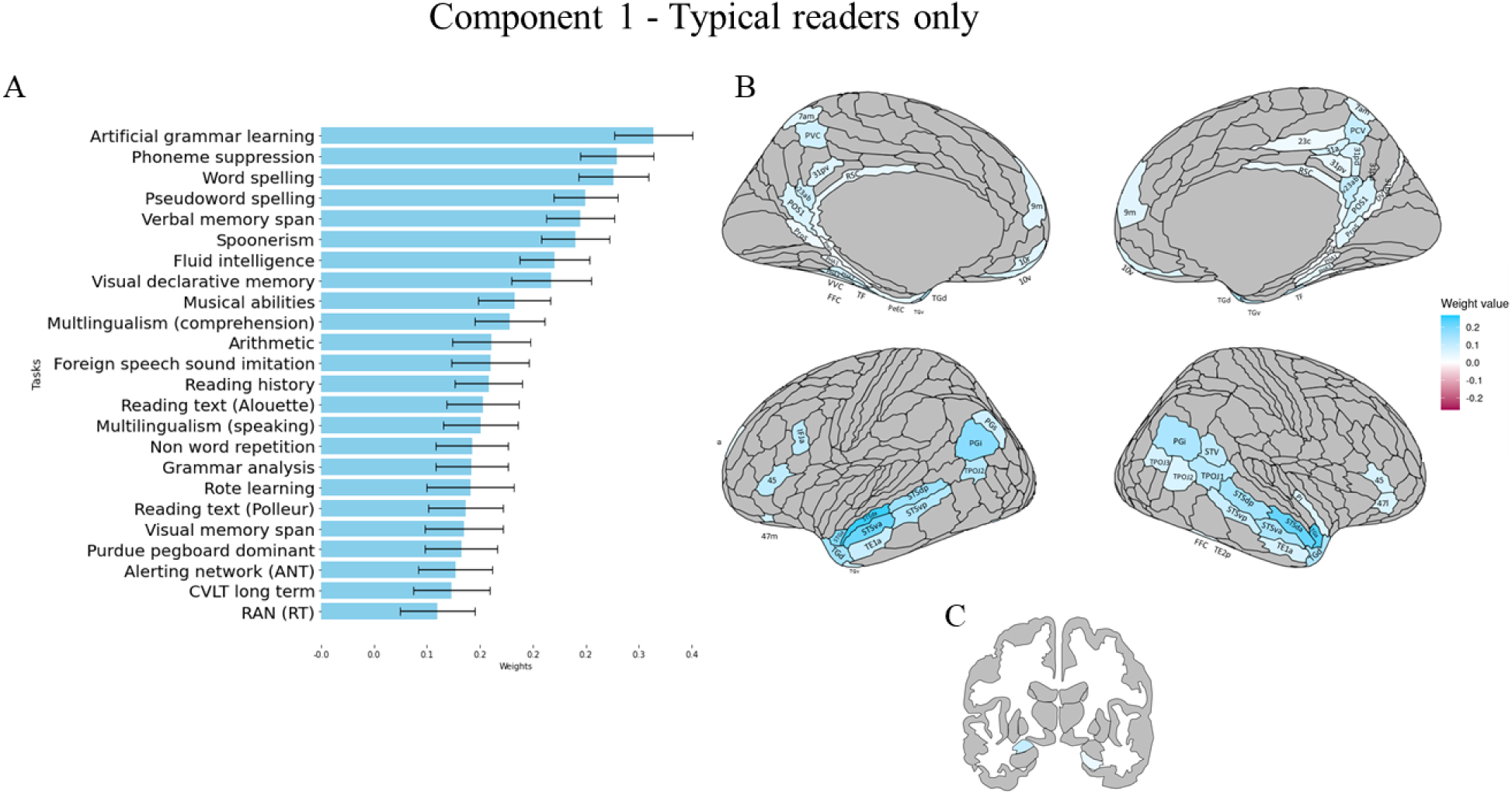
Results of the behavioural, cortical and subcortical saliences/weights for the first significant correlational PLS component in the typical readers. Blue represents significant positive weights, red represents significant negative weights, and grey represents non-significant ones. (A) Contribution of behavioural variables to the component (for all scores, higher values represent better performance). (B) Contribution of cortical ROIs to the component. (C) Contribution of subcortical areas to the component. For each component, mean weights, SD and brain uncorrected results can be found in the supplementary material, in the Supplementary results section.

A positive association was found with memory measures (such as Nonword repetition, Visual declarative and Verbal memory span, and Rote learning), pattern recognition and learning (Grammar analysis, Fluid intelligence, Artificial grammar learning), auditory measures including working memory: musicality and phonological awareness (including Spoonerisms, Nonword repetition, and Foreign speech sound imitation), and to a literacy measure involving sound-symbol correspondences (i.e. Word spelling). Positive associations were present also for multilingualism scores for comprehension and speaking, alerting skills and motor dexterity (Purdue pegboard test). Moreover, focusing on the differences in the present TRs analysis compared to that in the full sample, here component 1 was associated with additional measures for memory (CVLT long term, Visual memory span), arithmetic, literacy and literacy mediators (RAN, both Reading text measures, Reading history). Also, even though this component includes measures of phonological abilities in both the TRs and the whole-sample analyses, in this analysis including only the TRs group, the phonological measures play a more prominent role (i.e. higher weights). This suggests a stronger influence of phonology, which is particularly interesting when considering that this component also encompasses literacy measures. Finally, in the whole sample, multilingualism exposure contributes negatively to the component.

At the brain level, the ROIs contributing to the first component were similar to those contributing to the single component found in the whole sample, with more bilateral activation especially in temporal areas. Higher behavioural scores were associated with more activity in middle-to-superior posterior (STSdp, STSdp) and anterior (STSda, STSva, TE1a, STSva, STSda, STGa, TGd, TGV) temporal areas, in frontal areas (IFJa and 45 on the left, 45 and 47l on the right), in posterior inferior parietal regions (e.g. PGi, and surrounding clusters, again predominantly left-lateralised) and in temporoparietal regions (e.g. TPOJ2), the latter again somewhat right lateralised in extent. There was also a positive correlation with medial posterior cingulate (RSC, POS1, 23ab, 31pv and surrounding areas), and bilateral presubiculum (PreS) and parahippocampal (PHA1, PHA2, PHA3) areas. Subcortically, areas that contribute positively to the relationship are the right hippocampus and left amygdala, and the cerebellum on the right crus II. All results for cerebellar ROIs and uncorrected results can be found in the Supplementary Materials.

##### Component 2

The second component (Figure 4) accounted for 19% of the variance (*p* = .012, *r* = .35). This component was mostly negatively associated with tasks mainly related to motor and processing speed and lexical access compared to Component 1. Negative associations were present for motor speed, with Finger tapping speed (and not for dexterity, as in Component 1) and lexical access, reading and memory (RAN, CVLT short term, word and pseudoword reading, non-sensical text reading) and phonological perception. Two tasks measuring Visual declarative memory and phonological skills (Spoonerisms) were also positively associated with the component, with overall smaller weights compared to the negatively correlated measures.

**Figure 4.**
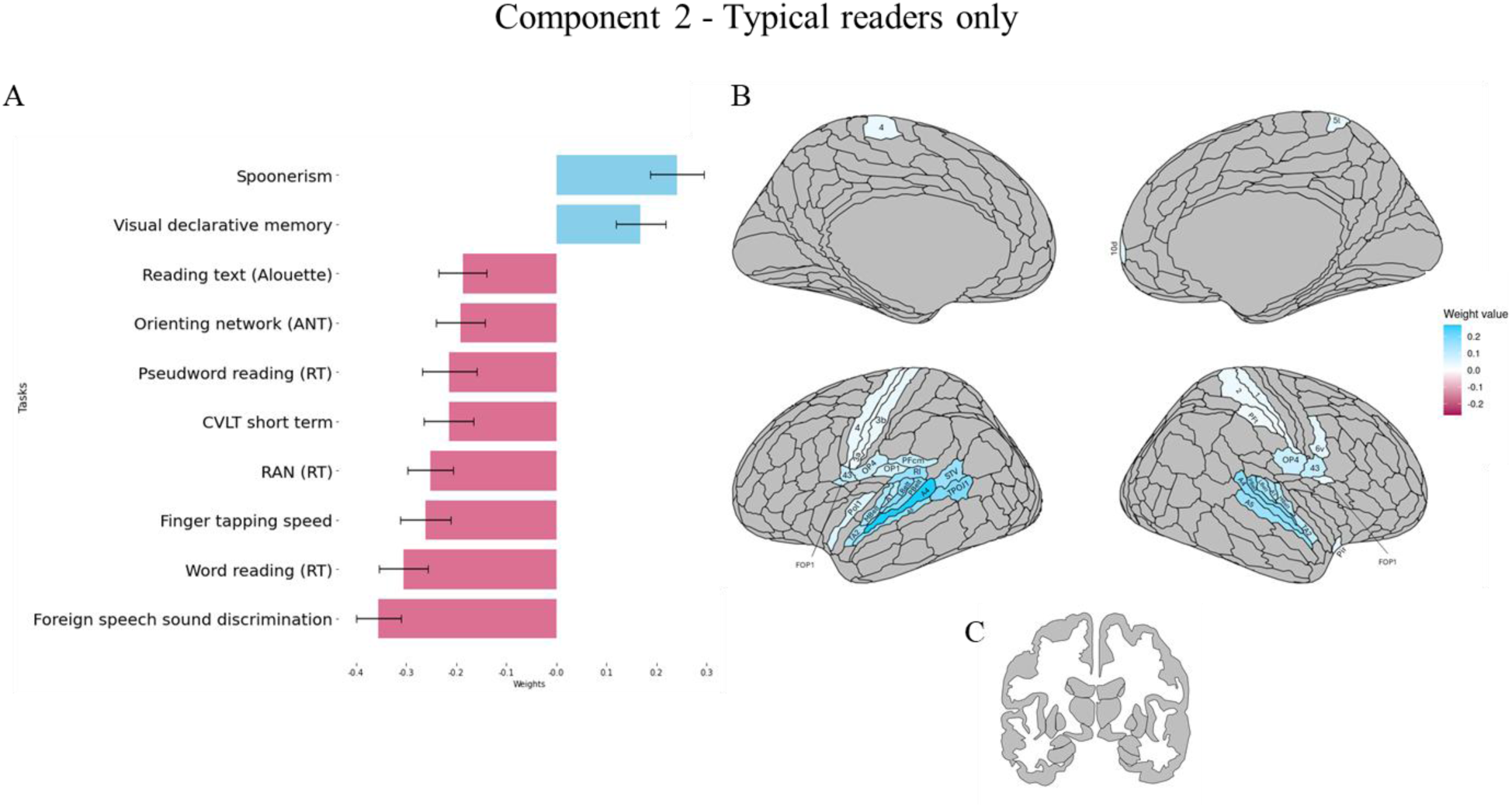
Results of the behavioural and cortical and subcortical saliences/weight for the second significant correlational component in the typical readers. Blue represents positive weights, red represents negative weights, and grey represent non-significant ones. Behavioural correlation for each individual variable to the component (for all scores higher values represent better performance). Cortical ROI correlations with the component. (C) Subcortical areas correlations with the component. For each component, mean weights, SD and brain uncorrected results can be found in the supplementary material, in the Supplementary results section.

At the brain level, mirroring the behavioural results, lower-level auditory and sensorimotor areas were associated with this second component. Poorer performance on the tasks negatively contributing to the component was associated with more activation in bilateral early auditory (A1, MBelt, PBelt, LBelt, A4, A5) and other auditory temporal association areas (TA2). Moreover, left primary motor (4) and bilateral sensorimotor cortices (areas 3a, 3b on the left, and areas 1, 2 on the right) were involved. Smaller positive contributions came from bilateral opercular/insular regions (e.g. OP4, 43, FOP1 bilaterally, OP1, RI, PFcm on the left). None of the subcortical or cerebellar ROIs contributed significantly to this second component. All results for cerebellar ROI and uncorrected results can be found in the Supplementary Materials.

##### Component 3

The third component accounted for 8% of the variance (*p* = .026, *r* = .67). This component showed the most complex pattern of results in the analysis. Figure 5A shows that it was behaviourally negatively associated with measures of auditory and phonological skill (e.g. Nonword repetition, Foreign speech sound discrimination, Phoneme suppression) and Verbal memory span score, and positively associated with a mix of cognitive, linguistic, and multilingual experience measures, some of which are negatively correlated with the previous components (e.g. Text reading, RAN, Orienting).

**Figure 5.**
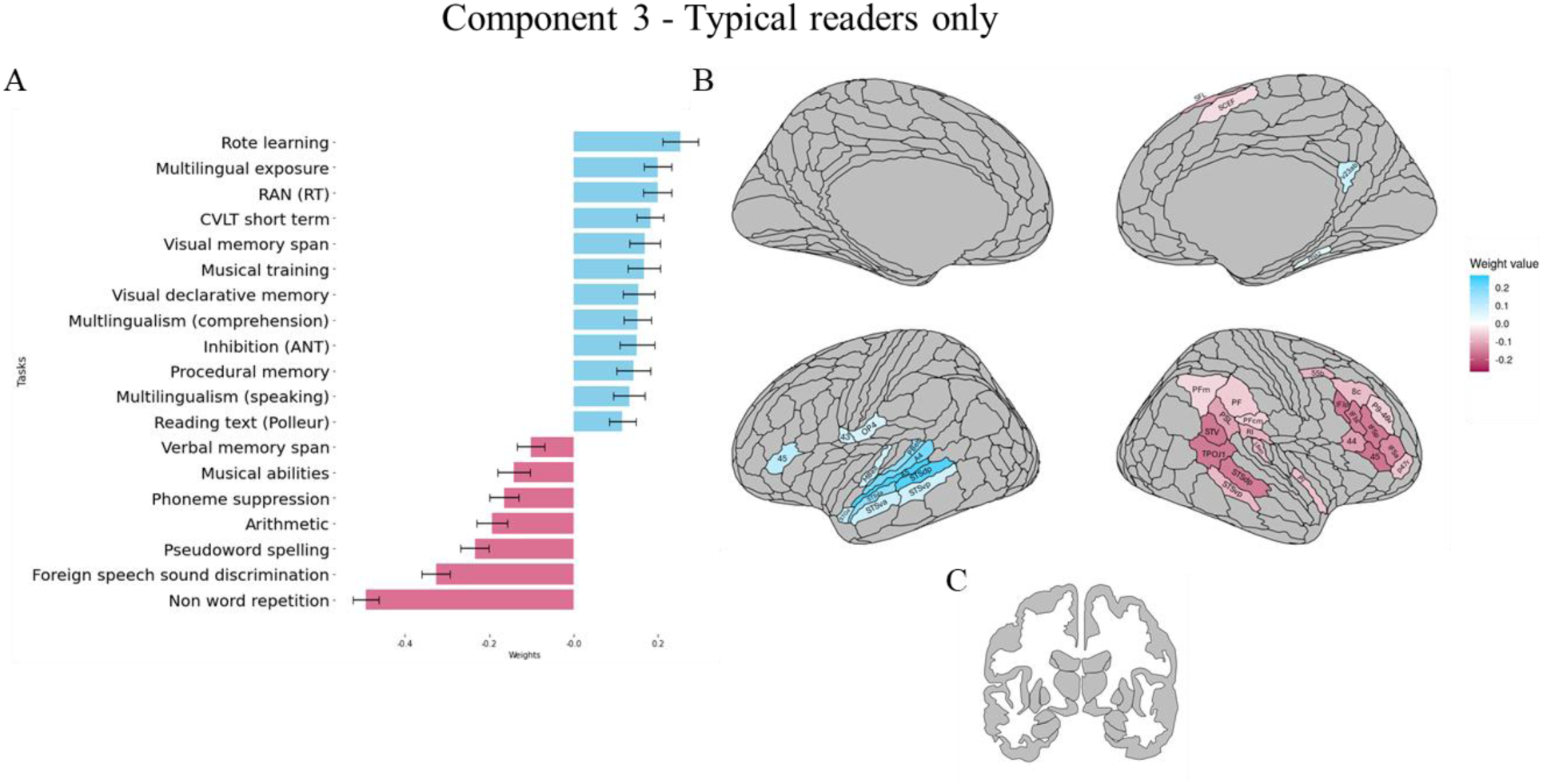
Results of the behavioural and cortical and subcortical saliences/weight for the second significant correlational component in the typical readers. Blue represents positive weights, red represents negative weights, and grey represents non-significant ones. (A) Behavioural correlation for each individual variable to the component (for all scores higher values represent better performance). (B) Cortical ROI correlations with the component. (C) Subcortical areas correlations with the component. For each component, mean weights, SD and brain uncorrected results can be found in the supplementary material, in the Supplementary results section.

In terms of relationships with brain activation, we observed a clearly lateralised pattern, mostly in higher-level language regions (Figure 5B). In the left hemisphere, regions contributing positively to the component included early auditory (MBelt, Pbelt) and associative auditory areas (STSda, STSva, STSdp, STSvp, STGa, A4, A5), as well as some clusters in opercular regions (45,43, OP4). Negative relationships were observed in right-lateralised areas such as the inferior frontal (IFJp, IFJa, IFSp, IFSa, 44, 45) and dorsolateral prefrontal cortices (55b, 8c, P9-46v, p47r), in the temporoparietal junction (e.g. TPOJ1, DTV), in the posterior STS and in other medial clusters (SFL, SCEF). In the right hemisphere, positive associations were only found with a left parahippocampal area (PHA2) and with one region in the posterior cingulate cortex (v23ab). None of the subcortical regions were significantly associated with this component (Figure 5c), and only the left cerebellar area VIIb negatively contributed, though marginally, to the component (Supplementary material).

## 4. Discussion

We investigated brain-behaviour associations by examining, in a data-driven and multivariate manner, behavioural measures of linguistic, domain-general and domain-specific skills and brain activation during story listening in the native language. We performed two PLS analyses, one on the full sample including TRs and dyslexic DRs readers, and the other only in typical readers. The former analysis allowed us to explore brain-behaviour relationships across a wider range of performance – one which included a (language/reading) learning disorder, and the latter allowed us to determine which relationships are more likely to be observed in typical readers. Results that are consistent across the two analyses are likely to reflect more stable and potentially more generalisable results. Results that differ between the two analyses are likely to reflect differences arising from the additional inclusion of individuals with a previous history of reading disorder, and/or differences arising from the inclusion of individuals with lower multilingual language experience, given that DRs on average has a lower multilingual language experience and exposure than the TRs.

In the next sections, we will discuss the brain-behaviour multivariate components arising from the two analyses, in light of how brain activation during speech processing is related to positive or negative relationships with subsets of tasks.

### 4.1. Overview of the components

#### 4.1.1. Component 1

Behaviourally, Component 1 (accounting for 36% of the brain-behavioural covariance in the full sample, and 44% of the variance in the TRs) in both samples showed an association between higher-level cognitive and linguistic skills, particularly for memory and grammar, and phonological awareness. These were associated with neural activation in higher-level areas within the language network known to be involved in combinatorial processes, in amodal semantic hubs, and in areas involved in phonological processing. In the TRs sample, behaviourally this component also included motor dexterity, and literacy measures, for text reading and spelling, with a heavier involvement of memory and phonological skills. This suggests that while language and cognitive measures are strongly and consistently associated across samples, literacy also becomes part of this association when only typically reading participants are included.

Overall, there are substantial similarities between the brain-behavioural association observed in the two analyses. In more detail, two of the tasks that positively contributed to this component reflect grammatical rule learning and knowledge (i.e. Artificial grammar learning and Grammar analysis) and pattern recognition (i.e. Fluid intelligence) skills, which reflect complex pattern analysis, arguably important components of cognition and intelligence. In fact, Artificial grammar learning was the strongest behavioural contributor to the brain-behavioural relationship. Neurally, activity in the bilateral dorsal and ventral anterior temporal lobes (ATL) was one of the strongest contributors to the brain-behaviour relationship of this component, again in both analyses. The ATL has been shown to be sensitive to abstract syntactic structure (Fedorenko et al. 2010; Pallier et al. 2011), and to be involved in lexico-semantic and compositional semantic processes (Hickok and Poeppel 2007). The left IFG (area 45) also contributed to this component; this area is known to support morphosyntactic processes (Hickok and Poeppel 2007) and is proposed to be involved in the semantic combinatorics (Friederici 2006; Schell et al. 2017).

There were also positive behavioural contributions of tasks that support or are related to lexico-semantic processing, such as memory and vocabulary learning tasks (e.g. Verbal memory, Rote learning and Visual declarative memory scores). Both brain imaging research and neuropsychological work on patients with stroke or neurodegeneration support the role of the ATL as an amodal, domain-general hub for semantic and conceptual knowledge and memory (Lambon Ralph et al. 2010; Lambon Ralph et al. 2016; Gainotti 2017). In both analyses, we found contributions of other brain areas that are also known to be amodal semantic hubs, where unimodal information converges: the bilateral IPL (PGi, PGs), bilateral posterior cingulate/retrosplenial (e.g. RSC, POS1, v23ab), medial prefrontal (9m, left-lateralised in the whole sample and bilateral for TRs), bilateral parahippocampus, and subiculum areas in the right hemisphere and entorhinal cortex in the left hemisphere (Fernandino et al., 2016).

Another set of tasks positively correlated with this component pertained to phonological/auditory processing and working memory (e.g. Foreign speech imitation, Spoonerisms, Phoneme suppression, Musical abilities). These tasks contributed in both analyses, but more so in the TRs analysis. These skills may partly drive the correlation with the activation in the posterior STS, known to be involved in speech and phonological processing (Hickok and Poeppel 2007; Hein and Knight 2008). Also, the relationship with phonological and language processing (imitation and multilingualism) may be driven by the activation of the IPL, a central hub in the dorsal phonological route of reading (Kuhl et al. 2020; Turker and Hartwigsen 2021). The IPL is also associated with sound-based phonemic representations (Coslett and Schwartz 2018), perception of ambiguous speech (Kilian-Hütten et al. 2011) and works in combination with IFG areas to support phonological memory processes (Kellmeyer et al. 2013).

The prominent role of grammatical processing and of verbal declarative memory in this first component inevitably recalls Ullman’s declarative/procedural model (Ullman 2004; Ullman 2016). While the model predicts that in L1, lexicon depends on the declarative memory system and grammar on the procedural one, it also posits that for L2, especially if explicitly learned and at low level of proficiency, both of these language components would depend on explicit and effortful retrieval from the declarative memory. The tasks associated with this component tapped into memory for lexical items and memory for verbal items, as well as into the learning and analysis of artificial languages, which are expected to assess L2 grammar learning processes. In line with this, our results show that procedural memory (measured through a serial reaction time task) did not contribute to the component. However, the declarative-based tasks that do contribute to the component correlate positively with brain activation in areas supporting declarative processes, in combination with a set of semantic and amodal conceptual hubs. This suggests that the learning and use of novel (i.e. L2-like) grammatical rules and verbal memory may rely on declarative skills and underlying medial temporal structures, as suggested by the declarative/procedural models, but that they are additionally supported by combinatorial and amodal semantic systems.

#### 4.1.2. Component 2

Component 2 (accounting for 19% of the brain-behavioural covariance), obtained in the analysis with TRs only, represents a complementary, lower-level component compared to Component 1. Behaviourally, there are negative contributions from tasks reflecting automatization, processing and motor and processing speed and lexical access (Penolazzi et al. 2007). What appears to commonly underlie most of the tasks showing negative weights is that most require fast processing and quick reactivity (RAN, Orienting and Finger tapping speed). In terms of the brain, lower-level auditory and sensorimotor cortices contribute positively to this component. The negative relationship between the brain and most of the behavioural measures of this component could reflect optimisation or efficiency, with lower behavioural scores being associated with increased brain activation. There were also, however, positive contributions of Spoonerisms, measuring phonological awareness, and Visual declarative memory to this component.

In more detail, bilateral activation of auditory and sensorimotor areas was associated with the above tasks, with a left-lateralised bias particularly in superior temporal regions and in the frontal opercular regions. Lower behavioural scores were associated with greater activation in auditory areas including the planum temporale, which are important for both phonological and orthographic processing in good readers (McNorgan et al. 2014). Moreover, greater activation was present in the TPJ, known to be important for speech-related auditory-motor association, this being a fundamental step in lexical access (Davis 2016 Jan 1) and being required in several of the tasks that are negatively associated with this component.

Most of these tasks also require speech production, which can explain the involvement of frontal opercular and motor areas. Bilaterally, the posterior frontal opercular/insular area 43 is associated with movements such as swallowing, area FOP1 is associated with imagined movements, and OP4 integrates sensorimotor information into actions (Baker et al. 2018).

There was also activation in the left motor (area 4) and sensorimotor areas (areas 3a and 3b). Areas 4 and 3b are thought to be involved in the control of hand/finger movements and in the processing of finger-specific information, respectively (Baker et al. 2018). The involvement of motor cortices is consistent with the correlation with our motor speed task. Moreover, left area 4 has been shown to be active during passage and alphabet writing tasks (Beeson et al. 2003).

#### 4.1.3. Component 3

Component 3 (accounting for 8% of the brain-behavioural covariance), again identified only in TRs, showed a more complex pattern of associations. Behaviourally, it was positively associated with cognitive and multilingualism scores, and negatively with auditory/phonological abilities. These performances were associated with more activation in left superior temporal and some inferior frontal/opercular areas, and with less activation in right hemispheric IPL/TPj and IFG/DLPFC areas.

In particular, better memory, inhibition and multilingualism scores were associated with lower activity in the right inferior frontal sulcus regions (IFS; IFSa, IFSp, IFJa, IFJa), right middle frontal gyrus (area 8, p9-46v), right IFG (44, 45, p47r), and IPL/TPJ regions (e.g., TPOJ1, STV, PFm, PSL). The right IFS has been linked to music, reasoning, attention (Ruland et al. 2022) and to rule implementation (Demanet et al. 2016). The right middle frontal gyrus has been shown to be involved in attention re-orienting (Japee et al. 2015), and its connectivity to be negatively correlated with numeracy (Koyama et al. 2017), which is also negatively associated in this component. Moreover, the right IPL/TPJ region supports executive functions (including working memory and inhibition) (Rothmayr et al. 2011; Igelström and Graziano 2017) and together with right IFG areas it is part of the ventral attention network, underlying bottom-up attentional processes (Corbetta and Shulman 2002; Igelström and Graziano 2017). At the same time, in this component, better scores on these tasks are also associated with poorer phonological awareness/musical abilities, and with greater engagement of left lateralized, early auditory (A5, A4, Pbelt, Mbelt), auditory association (STSda, STSvp, STSdp) and opercular areas (43, 45, OP4).

The relationships within this component suggest that increased multilingualism is associated with better short- and long-term memory for words and with better attention, but with poorer phonological awareness and poorer performance on tasks requiring working and short-term memory for sublexical (i.e. phoneme-level) information. This may be partly explained by the nature of our sample. Some of the most multilingual speakers (poly- and hyperpolyglots) in this study were highly proficient but non-native French speakers. Although the hyperpolyglots did not differ from the other participants in their behavioural scores on these tasks (as shown in the supplementary results, Table 10), the fact that they were performing tasks in a non-native language may have led some of them to rely more on lower-level auditory (e.g. acoustic) processes and on greater engagement of lower-level auditory regions to perform tasks requiring subtle phonological manipulations. Moreover, within this third component, higher multilingual experience is associated with clear brain functional lateralisation, suggesting greater engagement of a left-lateralised higher-level auditory (i.e. belt and parabelt regions including STS subregions and inferior frontal (i.e. BA 45) language network, and lower (maybe optimised) engagement of right-lateralised attention and executive areas. It should be noted, however, that this third (and last) component explains a smaller percentage of the variance (8%) compared to the other two (63% in total), and that therefore these results and interpretations may not be representative of broader and generalisable trends

### 4.2. Language and its components

#### 4.2.1. Language and cognition

Our data-driven analysis did not explicitly confirm or refute any of the traditional language models presented in this paper, nor was it our goal to compare language models. We aimed at more broadly exploring the relationship between performance on linguistic and non-linguistic tasks thought to be involved in aspects of language, and brain activation during a story listening task. Our results identified components that span most of the major areas considered essential for language processing, as well as additional regions.

In both samples, we have been able to show that, at the behavioural level, cognition and language present a stable association, suggesting a consistent overlap. This is in line with previous behavioural analysis, with different statistical methods, in this sample (Rampinini et al. 2024), and also in children tested with tasks tapping into similar domains (Udry and Berthele 2021). In both of our analyses, measures including memory and grammar learning abilities were strong drivers of the brain-behavioural association in the first, most important component. Previous work has successfully shown dissociations between language-specific and cognitive processes at both the brain and behavioural levels using finer-grained, targeted analyses, and areas responsive to language have previously been shown to be less responsive to domain-general cognitive tasks (Fedorenko and Varley 2016; Fedorenko et al. 2024). However, language does not occur in a vacuum. Our analysis involved exploration of the relationship between behavioural scores and *language-specific* activation rather than with task-free neural data (e.g. resting-state functional connectivity or brain structural metrics). As such, our results strengthen the case for the connection between language and cognition, since testing for associations with language-specific activation during story listening could have biased results in favour of language-specific tasks and/or of brain activation underlying language subcomponents (i.e., phonology vs. lexicon/semantics), while minimizing the role of cognitive measures. However, a consistent association between language and cognition was observed in the brain-behaviour components, despite the use of brain activation for story listening in the analysis which could have favoured language-specific outcome.

As discussed in the introduction, the overlap between language-specific processing and other domain-general and domain-specific abilities is predicted in light of evolutionary and developmental mechanisms and the nature of language in interaction. Language is mostly used in interaction with other individuals (Levinson and Holler 2014), and the use of language in real-life contexts requires us to engage in other cognitive processes beyond simply retrieving semantic and syntactic information, in order to keep track of the narrative, turn-taking and conversation (Hasson et al. 2018). Moreover, language and cognitive processes mutually influence each other during language learning (Kidd et al. 2018; Kidd and Donnelly 2020; Schwering and MacDonald 2020; Donnelly and Kidd 2021). The principles of Hebbian plasticity (i.e., “what fires together, wires together”) suggest that, at the neural level, interactions between the brain functional underpinnings of different processes can establish enduring functional connections between them. It is likely that the interaction between these skills during development, language learning and language use results in long-lasting connections between these domains throughout life, both at behavioural and neural levels (Munakata and Pfaffly 2004; Caporale and Dan 2008). Additional support for this idea comes from clinical work, where neurodegeneration has been observed to spread along established brain networks (Bak and Chandran 2012; Zhou et al. 2012; Drzezga 2018). One example of this is found in Motor Neuron Disease, in which the linguistic skills that are more impaired are those with the closest brain functional links to the motor system, with earlier impairment of verb and action processing than object naming (Bak and Chandran 2012). Taken together, these findings support the view that language and cognition are deeply intertwined at both behavioural and neural levels, shaped by shared developmental trajectories, overlapping neural circuitry, and continuous interaction throughout life.

#### 4.2.2. Complementary patterns for language and cognition

Examining the components obtained in the TRs reveals a pattern of complementarity, particularly between the components explaining most of the variance, Components 1 and 2.

Component 1 was predominantly linked to higher-order cognitive and linguistic skills, as well as domain-specific skills, and literacy measures such as spelling and text reading. It is known that higher instructional time with opaque orthographies is associated with better performance on spelling (Zaretsky et al. 2009). Higher instructional time with opaque orthographies is also related to text reading fluency, which requires additional skills beyond the necessary automatized ability to read single words, such as sequential processing of words (van Viersen et al. 2024). This could potentially indicate that this component is linked to higher-level, well-established skills. Even in the motor domain this component was more strongly associated with more complex motor abilities, like dexterity and fine motor coordination as measured by the Purdue Pegboard Test, rather than simple motor speed, assessed through speeded tapping. This component was reflected in the cortical activation of higher-order language and cognitive areas, where increased activation correlates with better behavioural performance. This suggests an active role of these regions during language processing, particularly in individuals with automatised, proficient reading skills and more multilingual experience.

Conversely, the second component captured a subset of lower-level abilities, including motor speed, rapid lexical access, and reading single items and nonsensical text. This component was mostly associated with activity in sensory and motor cortices. Higher activation in these regions correlated with poorer behavioural performance, suggesting that people with lower abilities rely more heavily on lower-level cortical regions.

This component appears to show a complementary pattern in terms of the behavioural tasks involved, with Component 1 involving higher-level, well-established skills and Component 2 involving lower-level, speed-related skills. Furthermore, at the neural level, Component 1 is primarily associated with higher-order association cortices, which are predominantly transmodal, whereas Component 2 is primarily associated with lower-order sensorimotor cortices, which are typically considered to be unimodal (Sydnor et al. 2021). The complementarity between the 2 components is likely to reflect an interplay between the two key aspects that characterise a typical skilled reader. Individuals with stronger linguistic, cognitive, musical and higher-level reading skills and with greater multilingual experience appear to use higher-level language and cognitive brain areas more actively, while relying less on lower-level sensorimotor regions.

#### 4.2.3. Literacy and other language functions

The association between language/cognitive scores and literacy may be unexpected given the previous work by our group (Rampinini et al. 2024) in which both in the full sample and in TRs only, literacy measures tended to consistently cluster separately from other language/cognitive scores. However, we need to consider that the analysis methods were different across studies, and that here, we explored the relationship between different data types, i.e. brain and behaviour, and not just between variables within one modality. We must consider that “cognitive processes and neural processes are not the same thing” (Valian, 2016, p. 570), and while literacy tasks can be dissociated from other language/cognitive skills at the behavioural level, they could rely on mostly shared underlying neural substrates.

In the present study, the differences between the analysis in the full sample (i.e. including the DRs) and the analysis including only the TRs showed that in the latter, these cognitive and linguistic processes are also associated with measures of literacy, particularly for reading fluency and history, while also showing a stronger association with phonological awareness (accompanied by more STG activation). This may suggest that in typical readers, literacy skills may rely on, or be more strongly linked to, more general cognitive/linguistic abilities and to their associated high-level neural mechanisms, reflecting a more specialised and expert, reading-ready cognitive architecture. On the other hand, in dyslexic readers, these literacy tasks engage different brain areas (i.e. not those language / cognitive ones associated with Component 1), perhaps as compensatory or more variable (e.g. noisy; Hancock et al., 2017) neural activations. Moreover, it suggests that when TRs only are considered, spoken language and literacy skills (along with phonological mediators) are associated to overlapping neural substrates. This finding is in line with the print-speech convergence hypothesis, which postulates that successful reading is supported by strong overlap between the neural substrates underlying the processing of written and of spoken language (Shankweiler et al. 2008; Rueckl et al. 2015; Preston et al. 2016).

#### 4.2.4. Multilingualism

In this study, we characterised multilingualism using 3 separate entropy scores that continuously captured participants’ language experience in terms of proficiency, comprehension abilities and current exposure. Importantly, multilingualism was not treated as an independent variable or used to categorise participants into binary groups such as ‘monolinguals’ and ‘bilinguals’. While this approach does not allow us to draw strong conclusions regarding the differences in the components based on language background, we consider that our approach offers several advantages: (1) it provides more nuanced and ecologically valid measures of multilingualism compared to strict dichotomies (e.g., mono- *versus* bilingual), and (2) it allows for the exploration of brain-behavioural associations across the full spectrum of different aspects of the multilingual experience (i.e. proficiency, comprehension abilities and exposure).

In terms of the role of the multilingualism variables in the obtained components, multilingualism exposure contributed negatively to the first component in the whole sample only. This negative contribution may be due to the differences between the three multilingualism variables being measured. Multilingualism *exposure* assessed the number of languages currently present in the participants’ environment, whereas the other two scores — multilingualism *speaking* and multilingualism *comprehension* — better capture language knowledge. The participants showed considerable variation in these latter 2 measures, with some polyglots speaking or understanding more than 10 languages. As a result, the current exposure measure may be less informative than our two measures of multilingual knowledge, as polyglots might not have been exposed daily to all the languages they know. In contrast, less multilingual participants (amongst which we can find DRs) were likely exposed to their known languages daily due to the multilingual nature of the region where data collection took place. Thus, while multilingual speaking and multilingual comprehension scores are strongly positively associated, multilingual exposure represents an anomaly. We may have very multilingual speakers with relatively low current exposure, making this measure less informative. This could explain why only language knowledge, represented by measures assessing cumulative experience with speaking and understanding one’s languages, were positively related to linguistic and cognitive skills. Conversely, current exposure to language, which provides a snapshot of the language environment at the time of assessment, did not show the same relationship. Once DRs were removed, multilingual exposure remained negatively associated with the component but not significantly, possibly indicating the reduced variability between these three measures given that DRs are less multilingual.

It is important to note that the participants in this study were, on average, highly multilingual, with very few monolinguals and bilinguals. Future works is necessary to confirm that our results are reproduced in the lower levels of the multilingual distribution.

## 5. CONCLUSIONS

In this paper, brain-behaviour associations for language-relevant skills and speech processing were investigated using PLS correlation analysis. Participants with different levels of multilingual experience and reading ability were included, and the analyses were performed first on the whole sample and then on the subset of TRs only, to explore which associations were stable and which were modulated by the presence or absence of DRs.

Overall, in both samples, a main component including cognitive, linguistic and phonological measures was found to be associated with cortical areas involved in lexico/semantics, combinatorial processing and amodal conceptual-semantic areas. Analyses excluding DRs suggest that literacy and phonological processes may be more closely associated with language and cognition in TRs than when DRs are included in the sample, pointing to more shared neural resources between these processes in TRs. Furthermore, results from TRs only indicate that additional components can be identified in populations without reading deficits, in particular, a complementary component involving foreign speech processing and motor speed, automatization and lexical access related to auditory and motor brain areas.

This work is a first step in exploring complex relationships underlying language use and learning and domain-general and domain-specific non-linguistic skills, in a multivariate way. We believe that language, as a multifaceted and multicomponent ability, should be studied comprehensively, that it should include broad behavioural characterisation of speakers and that it should ideally explore the relationship between different data modalities (e.g. brain function and structure, functional and structural connectivity, genetics, etc.). We believe that this approach can advance our understanding of language, of how individual differences arise and of how different abilities interact, and hope that future work will continue to unveil new aspects of these complex relationships.

## 6. ACKNOWLEDGEMENTS

We are very grateful for the statistical and methodological support for the PLS analysis offered by Erik Ringen, Farnaz Delavari, and Maria Giulia Preti. We would like to thank Michael Erard, Richard Simcott, Alexandre Teiga, Susan Fitzgerald, Lane Greene, Hugues Pluvinage and Swiss Radiotelevision for their help in the recruitment of the participants in general and of the polyglots in this study. We also thank Neiloufar Family and Sevil Maghsadhagh for rating our foreign speech sound imitation task.

## 7. AUTHORS’ CONTRIBUTIONS

Conceptualisation, I.B., A.R., O. K, R. B., N.G.; Data curation and preprocessing, I.B., A.R. and O.K.; Methodology, I.B.; Visualisation, I.B., O.K.; writing—original draft preparation, I. B.; writing—review & editing, I.B., A.R., O. K, R. B., N.G, funding acquisition, R.B. and N.G.

## 8. FUNDING

This work was supported by the Swiss National Science Foundation [Grant #100014_182381], and by the NCCR Evolving Language, Swiss National Science Foundation [Agreement #51NF40_180888].

## 9. SUPPLEMENTARY MATERIALS

Supplementary materials are openly available at the project’s Open Science Framework page at https://osf.io/4zcn2/?view_only=f1f5f11ba91c4e168e5321a198b4dc7a

## Notes

### Competing Interest Statement

The authors have declared no competing interest.

### Summary of Updates

This version addresses reviewer comments, particularly regarding the theoretical framework. The introduction and part of the conclusion has been updated to reflect these changes

